# Diet influences resource allocation in chemical defence but not melanin synthesis in an aposematic moth

**DOI:** 10.1101/2023.02.24.529866

**Authors:** Cristina Ottocento, Bibiana Rojas, Emily Burdfield-Steel, Miriam Furlanetto, Ossi Nokelainen, Sandra Winters, Johanna Mappes

## Abstract

For animals that synthesise their chemical compounds *de novo*, resources, particularly proteins, can influence investment in chemical defences and nitrogen-based wing colouration such as melanin. Competing for the same resources often leads to trade-offs in resource allocation. We manipulated protein availability in the larval diet of the wood tiger moth, *Arctia plantaginis*, to test how early life resource availability influences relevant life history traits, melanin production, and chemical defences. We expected higher dietary protein to result in more effective chemical defences and a higher amount of melanin in the wings. According to the resource allocation hypothesis, we also expected individuals with less melanin to have more resources to allocate to chemical defences. We found that protein-deprived moths had a slower larval development, and their chemical defences were less unpalatable for bird predators, but the expression of melanin in their wings did not differ from that of moths raised on a high-protein diet. The amount of melanin in the wings, however, unexpectedly correlated positively with chemical defences, irrespective of the diet they were raised on. Our findings demonstrate that the resources available in early life have an important role in the efficacy of chemical defences, but melanin-based warning colours are less sensitive to resource variability than other fitness-related traits.

## 1. Introduction

Organisms need to invest simultaneously in various life-history traits such as growth, energy maintenance, reproduction, predator avoidance and protection from pathogens (Stearns, 1992; Roff & Fairbairn, 2007). According to life-history theory, limited resources need to be distributed among various competing traits, inevitably leading to trade-offs in resource allocation (i.e., resource-allocation hypothesis; Bazzaz et al., 1987; Stearns, 1989; Glazier, 2002). Examples include trade-offs between larval and adult traits (Stevens et al., 1999), or between predator-avoidance and immune function (Rigby & Jokela, 2000). In general, allocation costs lead to trade-offs between traits that shape the demography of populations (McCauley et al.,1990; Boggs, 1992) and can significantly impact eco-evolutionary dynamics in nature (Schroderus et al. 2010; Farahpour et al., 2018). Testing the effect of only a few (one or two) traits on the resource allocation strategy may overlook critical information about the costs, especially if those traits are prioritised by the organism for fitness investment in a specific life stage (Lindstedt et al., 2020; de Jong, 1993). In organisms with complex life cycles, such as insects, considering how resources are allocated across and within life stages is essential to understanding how trade-offs shape fitness and different traits (Burdfield-Steel et al., 2019; Lindstedt et al., 2016).

Melanin plays an important role in the warning signals of many aposematic organisms (Fabricant et al., 2013; Lindstedt et al. 2020), those in which a primary defence (e.g., a warning signal such as colour, odour, sound) is coupled with secondary defences (chemical, morphological, behavioural) (Poulton 1890). Melanin is a phenolic biopolymer widely present across the animal kingdom, whose function in organisms as distant as mammals and insects is similar despite differences in the enzymes and substrates that regulate its production across species (Sugumaran, 2002). In vertebrates, including humans, melanin has a pivotal role in protection against UV radiation (Jablonski & Chaplin 2017; Nicolaï et al. 2020), as well as in sexual and protective colouration (Cuthill et al. 2017). In insects, specifically, melanin is also crucial in physiological and ecological processes such as thermoregulation (True, 2003; Trullas et al., 2007; Lindstedt et al. 2009), immune defence (Wilson et al., 2001), wound healing (Bilandžija et al. 2017) and protection from predators (Hegna et al. 2013; Majerus, 1998). The expression of melanin is often genetically based (Ellers and Boggs, 2002; Lindstedt et al. 2009; van’t Hof et al. 2019) and tightly related to protein resources (Lee, Simpson & Wilson, 2008). Thus, the proteins necessary for the synthesis of melanin pigments can constrain the expression of life-history traits and chemical defences (Lindstedt et al., 2020; Galarza 2021), making aposematic signals that require melanin good candidates for testing the resource-allocation hypothesis.

To date, several studies have focused on testing how the production of secondary defences may imply trade-offs in the allocation of resources needed for the primary defences, but experimental studies testing both the melanin component of warning signals and the chemical defences in the same system are lacking. A study on the Asian lady beetle, *Harmonia axyridis*, found no phenotypic correlation between the reflex bleeding response (frequency of secretion of the defensive chemicals) and the beetle colouration, even though reflex bleeding is costly and affects life-history traits (Grill and Moore, 1998). Recent work on monarch butterflies, *Danaus plexippus*., showed a link between toxin sequestration and warning signals, where male conspicuousness was inversely correlated with oxidative damage (due to an increase in concentrations of sequestered cardenolides) (Blount et al., 2023). Previous work by Lindstedt et al. (2010) showed that the aposematic, generalist herbivore *Arctia plantaginis* (hereafter referred to as the wood tiger moth) develops a paler, orange warning signal when reared on *Plantago lanceolata* with high concentrations of iridoid glycosides (IG) in comparison to the more conspicuous, dark red warning signal of individuals fed on a strain of the same plant with low IG concentrations. This suggests that the cost of conspicuousness arose via higher excretion costs rather than via resource-allocation costs (Lindstedt et al. 2010).

Here, we use the aposematic wood tiger moth, which has a black melanin pattern that covers approximately 20 - 70 % of its hindwings and 50 - 80% of its forewings (Hegna et al., 2013), to investigate how early-life resource availability influences melanin expression in the wings, the efficacy of chemical defences, and life history traits. In this species, melanin synthesis competes against the production of chemical defences for energy and resources from diet precursors (nitrogen from proteins). While the metabolic pathway in the wood tiger moth chemical defence has not been identified yet, the thoracic defence fluid present two methoxypyrazines which are, similarly to the eumelanin components, heterocyclic compounds based on nitrogen (Higasio & Shoji, 2001). We manipulated the protein content of the diet of male wood tiger moths, to investigate how wild-caught, natural predators (blue tits, *Cyanistes caeruleus*) respond to their chemical defences, and whether the production of such defences leads to trade-offs with the production of other traits, such as wing melanisation. We predicted that, compared to males raised on a high-protein diet, males raised on a low-protein diet would have (1) higher melanisation, resulting from a trade-off between melanin production and the effectiveness of secondary defences (i.e. more melanised individuals have less deterrent chemical defences, and vice versa); (2) less effective chemical defences, rendering them more palatable to bird predators; and (3) higher life-history costs in terms of size or developmental time.

## 2. Materials & Methods

### Study species

The aposematic wood tiger moth *Arctia plantaginis*, formerly *Parasemia plantaginis* (Rönkä et al., 2016), displays a conspicuous colour polymorphism (Watson and Goodger, 1986; Chinery, 1993; Nokelainen, 2013) throughout the Holarctic region (Hegna, Galarza, Mappes, 2015). Male hindwings across the species distribution range can have a red, yellow or white background with black patterning that covers variable proportions (ca 20-70%) of the wing. White and yellow colourations are partly produced by pheomelanin, whereas black is a dopamine-derived eumelanin (Brien et al. 2022). Previous research has scored melanisation (the amount of eumelanin) of the hindwings as “plus” (with stripes, high melanin) or “minus” (without stripes, low melanin) (see Fig. 1), but this arbitrary categorisation does not necessarily represent the true variation, in which subtle differences are not detectable by the human eye. Long-term natural frequencies of male wood tiger moth melanin morphs in Estonia (classified by human eye) are about 47% high-melanin and 53% low-melanin (O. Nokelainen, unpublished data; see Fig. 1).

**Figure 1.**
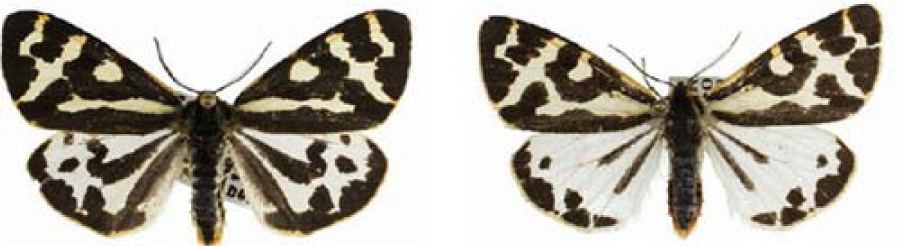
Hindwing and forewing melanin variation in male wood tiger moths from Estonia. Left, (+) high-melanin morph; right, (-) low-melanin morph. See text for details. Image from Nokelainen et al. 2013 (reprinted with permission).

As capital breeders, wood tiger moth adults do not feed; thus, resource acquisition only occurs during the larval stage. To avoid predation by avian and terrestrial predators, these moths produce two different types of defensive fluids: abdominal secretions deter predators such as ants, and thorax secretions deter birds with no adverse effects on ants (Rojas et al., 2017). The thoracic fluid contains two methoxypyrazines: 2-sec-butyl-3-methoxypyrazine (SBMP) and 2-isobutyl-3-methoxypyrazine (IBMP), which are produced *de novo* (Burdfield-Steel et al., 2018).

### Insects rearing, larval diet and thoracic fluid collection

Male wood tiger moths were obtained from a laboratory stock founded in 2013 with wild-caught individuals from Estonia and kept at the greenhouse of the University of Jyväskylä. The stock was supplemented annually with Estonian wild-caught individuals to preserve genetic diversity. The greenhouse conditions approximately followed the outdoor temperatures and natural daylight from May to August in Central Finland: between 20°C and 30°C during the day and decreased to 15°C-20°C during the night. We picked 10 families that contained both low- and high-melanin male morphs in the parental and grandparental generations, as classified by eye (see Fig. 1). Laboratory crosses show that wing melanisation (both pattern and amount) is strongly heritable (personal observation). After hatching, larvae of each of the 10 families were kept together for 14 days and fed with lettuce and dandelion (*Taraxacum spp.)*. Then, using a split family design, they were divided into two artificial diet treatments that differed in protein content (high protein, low protein; see recipe in supplementary material). For each diet treatment, we used 240 individuals which were kept in boxes of 10 individuals until reaching the pupal stage. Each box was checked and watered daily and cleaned when needed. Larvae were fed daily, *ad libitum*, with their corresponding diet treatment. Life-history traits (time to pupation, pupal weight) and the degree of melanin (high +, low -) were recorded for each individual. In total, 480 larvae were followed from eggs to adulthood or death, 177 of which emerged as females and 184 as males. All larvae were grown by Furlanetto (2017) from May to August 2017. When the male adults emerged from pupation, they were given water and stored at 4°C to slow their metabolic rate and to maintain their condition; thoracic chemical fluid was then collected in June and July 2017. Prior to fluid collection, moths were kept at 20-25°C for thirty minutes. Then fluid was extracted by squeezing just below the prothoracic section with tweezers and collecting expelled fluid in a 10 μl glass capillary. Fluid samples were stored in glass vials at −18°C (Furlanetto, 2017).

### Effect of diet manipulation on melanin

To measure variation in wing melanisation, we photographed the 53 male moths whose defensive fluids were used in the predator response assay. The degree of melanisation varies continuously and thus we measured the proportion of the hindwing that was melanised on a continuous scale using image analysis.

Spread male moths were photographed using an established protocol (Nokelainen et al. 2017) and scaled to the same resolution (pixels per mm). We used the MICA toolbox (Troscianko & Stevens 2015) in ImageJ (v. 1.50f) for image analysis. From every image, a set of regions of interest (ROIs) were selected from the dorsal side of the wings: (a) whole forewing area, (b) forewing melanised region, (c) whole hindwing area, and (d) hindwing melanised region. The relative proportion of the wings that were melanised was calculated by dividing the area of the melanised ROIs by the whole wing ROIs, for the forewings:[(forewing melanised region area) / (whole forewing area)]; and for the hindwings: [(hindwings melanised region area)/(whole hindwings area)].

### Effect of diet manipulation on the predator response to male’s defensive fluid

Blue tits (*Cyanistes caeruleus*) are generalist feeders that have a similar distribution as *Arctia plantaginis* in Europe and are known to attack this species. Blue tits are common in Finland and can be kept briefly in captivity for experiments (e.g., Rojas et al., 2017). For the predator response assay (*C. caeruleus; n* = 62) we used fluids from 53 male moths fed on high- and low-protein diets; baits soaked with water were offered to 9 birds as a positive control. The volume of wood tiger moth thoracic fluid has no effect on blue tit responses (Burdfield-Steel et al. 2019), so we diluted each fluid sample with water to reach a total volume of 15 μl. The 15 µl was then divided into two samples of 7 μl each, which were used as bait for the same bird. The same amount of water was offered to control birds. The birds used for the experiment were caught at Konnevesi Research Station (Central Finland) from February to April 2018 using feeders with peanuts as bait (Ham et al. 2006), and then housed individually in plywood cages for the duration of the assays (see Ottocento et al. 2022).

Birds were first familiarised with the experimental boxes and trained to eat a bait (oat flakes) (see Ottocento et al. 2022). Each bird then experienced four sequential trials, at 5-minute intervals, in which they were presented with a plate containing one bait (see Fig. 2 A). The first and last trials were done with baits soaked in tap water to ensure that the bird was motivated to eat (first) and that the bird was still hungry (last). In the second and third trials, the bird was presented with a bait soaked in 7μl of the defensive fluids of the same moth (30 raised under a high protein diet, 23 raised under a low protein diet). The trial ended two minutes after the bird had eaten the whole bait, or after a maximum duration of 5 minutes if the bird did not eat the whole bait. In each trial, we recorded the proportion of the bait eaten (see Fig. 2 B), the beak cleaning frequency (number of times the bird wiped its beak against a surface, e.g., the perch), the latency to approach (how long it took for the bird to get close to the plate on which the bait was offered), the latency to eat after approaching (the hesitation time between approach and attacking) and the latency to eat.

**Figure 2.**
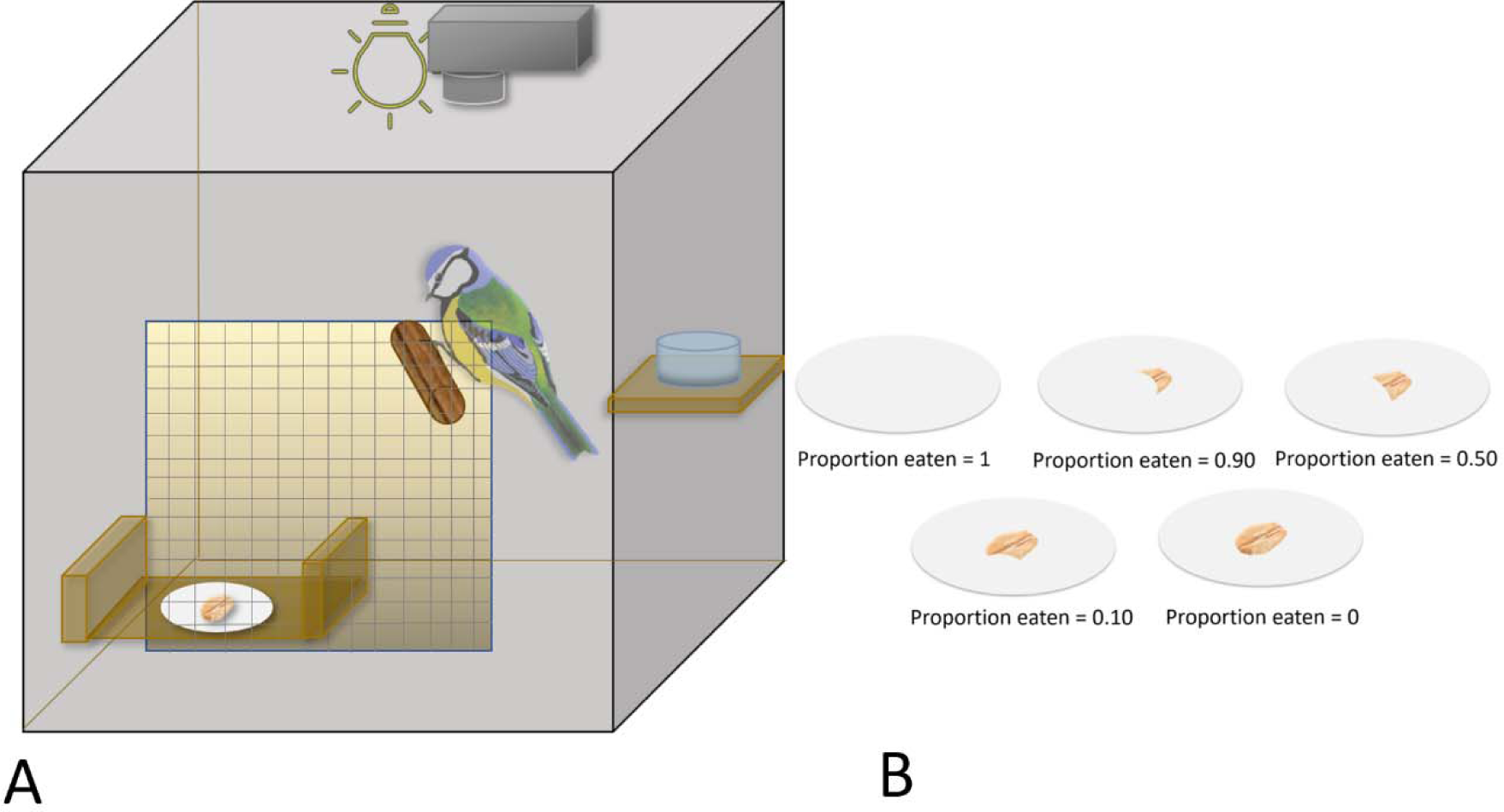
**A).** Experimental setup of the predator assay illustrating the perch, water, camera, light source, mesh opening for observation and hatch for inserting the plate with the bait (oat flake) into the enclosure. **2 B)** Proportion of bait (oat soaked with defensive fluid or water) eaten by the predators. The palatability was classified into five classes according to the proportion of the bait consumed by the birds: 100% (1), 90% (0.90), 50% (0.50), 10% (0.10), and 0%.

## Statistical analyses

All statistical analyses were done with the software R v. 4.1.2 (R Core Team, 2022) using the RStudio v. 1.2.1335 interface (RStudio Team, 2019). The level of significance in all analyses was set at *p < 0.05*.

### Effect of diet manipulation on melanin

Our results show that the amount of melanin in the hindwings of wood tiger moths is best measured as a continuous variable (see supplementary material), so we analysed how protein content in the diet influences the proportion of the wing that is melanised and the size of the wings using a linear mixed models with treatment as a fixed factor and family as a random factor, using the lme4 package (Bates et al. 2015). The total area of the forewings, the total area of the hindwings, the amount of melanin in the forewings (melanised area forewing/total area forewing), and the amount of melanin in the hindwings (melanised area hindwing/total area hindwing) were set as response variables (in separate models); the different types of diet (high/low protein content) were set as predictors.

### Effect of diet manipulation on the predator response to male’s defensive fluid

To test the proportion of bait eaten, we used beta regression models using package glmmTMB (Brooks et al., 2017) with family = beta_family(link= “logit); diet (low protein, high protein), the amount of melanin, and trial set as fixed effects; bird ID as a random effect; and an offset included to account for differences in observation time. To test for differences in beak wiping events per minute, we used generalised linear mixed-effects models (GLMM; Knudson et al., 2022) with a log link and Poisson distribution, fit by maximum likelihood (Laplace approximation). The interaction between diet (low protein, high protein), the amount of melanin, and trial were set as fixed effects, beak wiping frequency was set as the response variable, and bird ID was set as a random effect. To test the latency to approach, the latency to eat after approaching and the latency to eat, we used a Cox mixed-effects model using package *coxme* (Therneau, 2020), fit by maximum likelihood. These three behaviours were set as response variables; the interaction between diet (low protein, high protein), the amount of melanin and trial were set as fixed effects; and bird ID was set as a random effect. The behaviours of the predators were first compared to a water-only control to determine whether moth chemical defences elicited adverse predator reactions. We used Spearman’s rank correlations (Ojala et al., 2005) to test the correlation between the amount of melanin and predator responses to chemical defences (proportion of bait eaten, beak wiping, latency to approach, latency to eat after approaching, latency to eat).

### Effect of diet manipulation and hindwing melanin on life-history traits

To test whether male moths’ development time is affected by dietary protein and the amount of melanin, we used a mixed-effects Cox model (package coxme, Therneau et al., 2020), with development time included as the response variable, diet treatment and melanisation included as the predictor variables, and family as a random factor. We assessed the effect of diet treatment on pupae weight with a generalised mixed-effects model (package lme4, Bates et al. 2015) with a Gamma (link=“log”) distribution, with diet treatment and melanisation as the predictor variables and family as a random factor. We used a linear mixed-effects model (package lme4, Bates et al. 2015) with diet and melanisation as predictor variables and family as a random factor to test the effect of diet on the volume of the thoracic fluid. We tested the correlation between the pupal weight and the volume of the thoracic fluid using a Pearson correlation. Spearman’s rank correlations were used to test the relationships between the amount of melanin and the developmental time, pupal weight, and volume of thoracic fluid.

## Results

### Effect of diet manipulation on melanin

In moths raised in low-protein diet, neither the wing melanin content of the hindwings (coef ± s.e =0.001± 0.32, t = 0.265, *p = 0.96*) and the forewings (coef ± s.e = −0.02± 0.03, t = −0.85, *p = 0.40*) nor the size of the hindwings (coef ± s.e = −95528± 81245, t = −1.18, *p = 0.246*) or forewings (coef ± s.e = −104081±89682, z = −1.16, *p = 0.25*) differed to moths raised on a high-protein diet.

### Effect of diet manipulation and hindwing melanin on the predator response to male’s defensive fluid

To test if the thoracic fluid of the moth evoked an adverse reaction in the birds, we first examined predator behaviour in response to baits soaked in water (control). Baits soaked in the chemical defences of moths raised on a high-protein diet were eaten by the predators in lower proportions than baits soaked in water (control) (coef ± s.e −2.54 ± 0.87, z = −2.91, *p = 0.004*), while baits soaked in fluid from moths fed with low protein diet did not differ to the predators’ response to baits soaked in water (coef ± s.e −1.56 ± 0.87, z = −1.79, *p = 0.074*). We found no significant differences in the other predator behaviours recorded (i.e., latency to approach; latency to eat after approach; latency to eat; frequency of beak wiping) between birds exposed to baits soaked in either fluid from moths raised on high-protein or low-protein diets, and those exposed to water-soaked baits (*p > 0.05* for all comparisons, see Table S1A, S2A, S3, S4A; supplementary material).

Irrespective of diet type, the chemical defences of more melanised individuals were also more deterrent against predators (Spearman’s rank correlation, r_s_ = 0.27; *p = 0.005*, Fig. 3A), as baits soaked in these defences were eaten in lower proportions than those from less melanised males (coef ± s.e −5.20 ± 1.96, z = −2.66, *p = 0.007*, Fig. 3A). Similarly, the predators’ latency to eat the bait increased with higher amounts of melanin (Spearman’s rank correlation, r_s_ = 0.26; *p = 0.012*, Fig. 3B), indicating an adverse reaction of the predators to the chemical defence of highly melanised individuals, regardless of the diet they were raised on (coef ± s.e =-4.92 ± 2.17, z = - 2.26, *p = 0.024,* Fig. 3B). There was no correlation between the amount of melanin and other behavioural variables (latency to approach; latency to eat after approach; frequency of beak wiping; *p > 0.05* for all comparisons; supplementary material).

**Figure 3.**
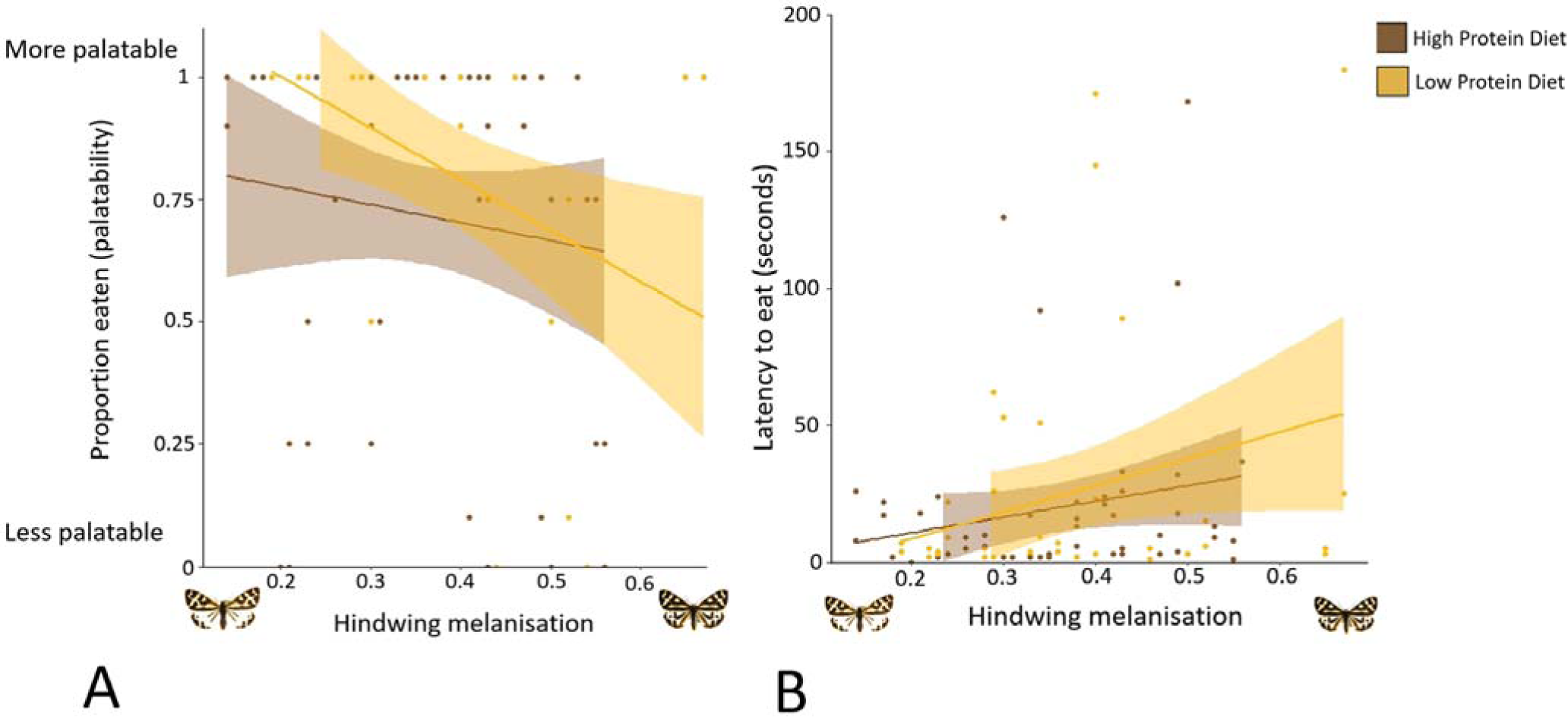
**A)** Palatability (proportion of defence fluid-soaked bait eaten by birds) and **B)** latency to eat the defence fluid-soaked bait compared to hindwing melanisation for male moths raised on high- and low-protein diets.

### Effects of diet manipulation and hindwing melanin on the life-history traits

The developmental time from egg to pupa varied among diet treatments: individuals raised on a low-protein diet took significantly longer to pupate than those raised on a high-protein diet (coef ± s.e =-0.92± 0.32, z = −2.87, *p = 0.004*). There were no differences in body mass (coef ± s.e= 0.03 ± 0.04, t = 0.75, *p = 0.47*, see Table S6) or in the volume of the thoracic defensive fluid (coef ± s.e =1.05± 0.87, t = 1.2, *p = 0.23*) between individuals raised on high- and low-protein diets. The volume of the chemical defence did not correlate with the pupae weight (t = 0.96, df = 188, *p = 0.34*). There was no correlation between the melanin amount and the larval developmental time (r_s_ = 0.11; *p = 0.435*), the body mass (r_s_ = 0.003; *p = 0.98*), or the volume of defensive fluid (r_s_ = −0.03; *p = 0.79*).

## Discussion

Investigating how resource allocation affects life-history traits and defences is vital for understanding which factors drive variation in the warning signals and chemical defences of aposematic organisms. Phenotypic variation in aposematic animals is puzzling because predators avoid defended prey that are common but attack rarer ones (Müller 1878). Trade-offs in resource allocation have been suggested to account for some of the variation found in natural aposematic populations (Wang, 2011; Burdfield-Steel et al., 2019; Blount et al., 2023). However, the phenotypic correlations among traits that characterise aposematic organisms such as melanin and chemical defences do not necessarily prove this connection, as interactions between such complex traits can be shaped by genetic correlations, pleiotropic effects or simply variable environmental conditions (e.g., food availability, environmental stochasticity, presence of predators). Therefore, to test the resource allocation hypothesis, we need to first test whether the production of defences is costly and then, whether their development is linked to other traits. To investigate whether those traits are costly, and whether a specific trait (melanin or chemical defence) is prioritised over others, we manipulated specific constituents of the early-life resources of male wood tiger moths. The diet used in the study varied in the amount of protein, which is an essential compound in different aspects of the larval life history of this species, and which has been shown to influence the efficacy of the species’ warning signal, immunity and life-history traits (Lindstedt et al., 2020).

Surprisingly, dietary protein levels had no effect on the amount of melanin in the fore- or hindwings, suggesting that this trait may not be directly influenced by early-life environment. However, Lindstedt et al. (2020) recently found that the protein content of the diet fed to larval wood tiger moths directly influenced the amount of forewing melanin of adult males in the closely related Finnish population. The apparent mismatch between Lindstedt et al.’s (2020) and our results could be due to the study design: Lindstedt et al. (2020) used larvae from selection lines for low and high larval melanisation, such that the phenotypic variation was larger than the natural variation (and, consequently, the average phenotypes were either rare or missing completely), likely making the possible costs of melanin easier to detect. In this study, by contrast, we used the natural variation of phenotypes. Interestingly, Lindstedt et al. did not find differences in the amount of hindwing melanin between individuals fed with high- and low-protein diets. Thus, hindwing melanisation and patterning seem to be under strong genetic control, which is unsurprising because moth hindwings commonly have an important signalling function (Sargent 1978, Kang et al. 2017; Rönkä et al. 2018). It is also possible that the Finnish and Estonian populations differ in their degree of plasticity of melanin production, but this hypothesis requires further investigation. Other studies in invertebrates have shown that melanisation is influenced by the amount of protein content in the diet (Lee et al., 2008; Ethier et al., 2015), while in vertebrates melanin is less affected by dietary changes (Hill and Brawner, 1998; Lee et al., 2008). For example, in the African cotton leafworm, *Spodoptera littoralis*, a diet with high protein content led to more melanised cuticles, faster growth and better antibacterial activity and survival (Lee et al., 2008). In the forest moth *Malacosoma disstria,* individuals raised under low-nitrogen availability had decreased melanic pigmentation and smaller size, highlighting the high costs of melanisation (Ethier et al., 2015). In contrast, in vertebrates such as the tawny owl, *Strix aluco*, the genetic control of melanin deposition appears to be strictly regulated (Roulin & Dijkstra, 2003; Mundy & Kelly, 2003; Bize et al., 2006; Hoekstra, 2006; Emaresi et al., 2011). However, even in vertebrates, this is not always the case and environmental conditions may alter melanised traits in some organisms (Fargallo et al., 2006).

The amount of melanin in the wood tiger moth wings increases with latitude and altitude (Hegna et al. 2013), as predicted by the thermal melanism hypothesis, which states that darker individuals have an advantage under low temperatures because they can warm up faster than light individuals (Trullas et al. 2007). Convincing support for the fitness advantage of dark colouration in cold environments has been found in many invertebrates (e.g., Kingsolver 1995; Ellers & Boggs 2004; but see Rosa and Saastamoinen 2020), but also in cold-blooded vertebrates (Vences et al. 2002; Stuart-Fox et al. 2017; Azócar et al. 2020). Interestingly, more melanin in the wings results in a better capacity to absorb radiation and warm-up for flight at the expense of increased vulnerability to predator attacks due to a reduction in the size of the non-melanic component of the warning signal (yellow or white pigment in the hindwings) (Hegna et al. 2013). Similar results were found also in wood tiger moth larvae, where higher amounts of melanin in the body resulted in improved thermoregulation at the cost of increased predation (Lindstedt, Lindström, & Mappes, 2008; 2009; Nielsen & Mappes, 2020). It is possible that the Estonian population, situated at lower latitudes than the Finnish population, is free from the (potentially genetic) constraints imposed by a colder climate. Moreover, while the melanin pattern in male wood tiger moths seems genetically regulated and the environmental component is low, our study shows that any dietary resources beyond what is required for the melanin patterns are allocated elsewhere, such as to more effective chemical defences.

We hypothesised that the thoracic fluid of males raised on a high-protein treatment would elicit a stronger predator response than the fluids from moths raised on a low-protein diet, as shown in a previous study (Furlanetto 2017) where the abundance of pyrazines was higher in fluids from moths raised on the high-protein artificial diet. An increase in protein content means an increase in the concentration of nitrogen, an essential element in the synthesis of the pyrazine molecule (Hodge, Mills & Fisher, 1972; Wong & Bernhard, 1988). Pyrazines have a characteristic repulsive odour (Rothschild, Moore & Brown, 1984; Guilford et al. 1987; Kaye et al. 1989; Moore, Brown & Rothschild, 1990) and a deterrent effect on birds (Marples & Roper, 1996; Lindström, Rowe & Guilford, 2001; Siddall & Marples, 2011). In this study, predators found the chemical secretions from male moths raised on the high protein diet more unpalatable (i.e., tasted worse) than those from males raised on the low protein diet, which confirms that the chemical defence is costly. However, we did not find any differences between the predators’ hesitation time (i.e., latency) to “attack” the bait soaked with fluids from male moths raised on high-protein and the latency to attack baits with water (control), which suggests that the predators may not perceive a difference in the odour (volatile compounds) of both. These results agree with previous studies suggesting that the defensive fluids of wood tiger moths may contain several repulsive compounds, not only pyrazines, that influence defence efficacy in concert (see also Winters et al. 2021; Ottocento et al. 2022) and affect predator response in different ways (Rojas et al. 2019; Winters et al. 2021). Moreover, as Ottocento et al. (2022) recently showed, the relative amount of the two methoxypyrazines (SBMP, IBMP) present in the wood tiger moth defensive fluids is more relevant to predator deterrence than the total amount of pyrazines.

Unfortunately, however, it is not possible to quantify the amount of pyrazines and conduct bird assays with the defensive fluids of the same individual. Interestingly, the same study (Ottocento et al., 2022) showed that predator responses are stronger towards the chemical defences of individuals originating from populations with high predation pressure (e.g., Scotland; Rönkä et al 2020) than towards individuals from populations with lower predation pressure (e.g., Estonia; Rönkä et al 2020), even if the total amount of pyrazines in the defensive fluids does not differ among populations.

Our results show that highly melanised male moths from both diet treatments have more effective chemical defences than those with less melanin, hinting at a positive correlation between costly melanin pigments and chemical defences. While this is not what we predicted, particularly in a scenario of low resource availability in early life, environmental conditions may have a greater influence on chemical defences than the pigmentary composition of the melanin hindwings, which is less sensitive to the variability of resources than other fitness-related traits. Although in this species the chemical defence compounds are not sequestered directly from plants but produced *de novo* (Burdfield-Steel et al., 2018), the main elements that the moths require to build their defences are still collected from their food intake.

When looking at how early-life resource availability influences wood tiger moths’ size, we hypothesised that males raised on a low-protein diet would have both smaller pupae and smaller hind- and forewings than those raised on a high-protein diet, as environmental fluctuations and environmental stress (in this case due to the low amount of protein in the diet) may strongly affect both insect wing (Bitner-Mathé & Klaczko, 1999) and pupal size (Nguyen et al., 2019).

Our findings, however, reveal no differences in size between males raised on high- and low-protein diets. This could be because individuals raised on a low-protein diet had a longer development time than those raised on a high-protein diet, which might facilitate the acquisition of more resources at the larval stage, even if of poorer quality, allowing these individuals to ultimately reach the same size as those on a richer diet, as reported by Lindstedt et al. (2017). A longer developmental time, however, may also lead to increased predation risk (Clancy & Price,1987; Häggström, & Larsson 1995), higher vulnerability to environmental perturbations (Tammaru et al., 2001), reduced fecundity (Saastamoinen, Hirai, & van Nouhuys, 2013), and difficulties to escape a risky environment (Cowan, Houde & Rose, 1996).

In sum, experimental evidence suggests that the production and maintenance of chemical defences are affected and limited by the resources available in early life, but melanin synthesis is not. This implies that, at least in wood tiger moths, melanin synthesis in adult wings seems to be less environmentally regulated than pyrazine production, implying that these traits are not limited by the same resource pool. Our findings also confirm that resource-dependent variation in chemical defences is perceived by natural predators, which has seldom been shown when variation in chemical defences is investigated (White and Umbers, 2021). Studying the response of natural predators, which are the selective agents for defended prey, is key when aiming to understand how variation in defences is maintained in aposematic species.

## Acknowledgements

We are grateful to Helinä Nisu, who assisted with the catching and training of wild birds. We thank Konnevesi Research Station for the use of the facilities. Kaisa Suisto and other greenhouse workers were invaluable in the rearing of the moths. We thank the members of the ‘Ecology and Evolution of Interactions’ group for the useful discussion on an early version of the manuscript.

## Funding

This study was funded by the Academy of Finland the project No. 345091 (to JM) and Societas pro Fauna et Flora Fennica (grant to CO).

## Author contributions

JM, BR and EB-S conceived and designed the study. MF, EB-S and BR raised the larvae under the different treatments. MF, ON, CO and SW did the colour analyses. CO, BR and EB-S did the unpalatability assays with birds. CO and BR wrote a first draft of the manuscript with input from JM and EB-S. All authors critically reviewed the manuscript, approved the submitted version, and agreed to be held accountable for the content therein.

## Data Accessibility

Helsinki repository

## Conflicts of Interest

None

## Permits

Wild birds were used with permission from the Central Finland Centre for Economic Development, Transport and Environment and licence from the National Animal Experiment Board (ESAVI/9114/ 04.10.07/2014) and the Central Finland Regional Environment Centre (VARELY/294/2015). All experimental birds were used according to the ASAB/ABS Guidelines for the treatment of animals in behavioural research and teaching (Association for the Study of Animal Behaviour, 2020).

## Notes

### Competing Interest Statement

The authors have declared no competing interest.

### Summary of Updates

August 16th, 2023: updated figure 2B "The palatability was classified into five classes according to the proportion of the bait consumed by the birds: 100% (1), 90% (0.90), 50% (0.50), 10% (0.10), and 0%. “The sample size of the total birds used (62) is now specified in the paragraph "Effect of diet manipulation on the predator response to males defensive fluid”

