## Supplementary_Material for "Diet influences resource allocation in chemical defence but not melanin synthesis in an aposematic moth" (from another version of this preprint: version 2, 2023-06-10)

**Address:**
[1]University of Helsinki, Organismal and Evolutionary Biology Research Programme, Faculty of Biological and Environmental Sciences, P.O. Box 65, Viikki Biocenter 3, Finland
[2]University of Jyväskylä, Department of Biology and Environmental Science, P.O. Box 35, 40014, Jyväskylä, Finland
[3]Department of Interdisciplinary Life Sciences, Konrad Lorenz Institute of Ethology, University of Veterinary Medicine Vienna, Savoyenstraße 1, 1160, Vienna, Austria
[4] University of Amsterdam, Institute for Biodiversity and Ecosystem Dynamics, Science Park 904, 1098 XH Amsterdam, The Netherlands

**Contents:** **Pages**

High and Low protein diet recipe 1

Is hindwing melanisation continuous or discrete? 2

Effect of diet manipulation and hindwing melanin on the predator response to male's defensive fluid 4

### 1. High and Low protein diet

| High protein artificial diet | Low protein artificial diet |
|---|---|
| Agar g 18 | Agar g 18 |
| Wheat Germ g 72 | Wheat Germ g 72 |
| **Teklad VitFree Casein g 32.4** | **Teklad VitFree Casein g 10.8** |
| Sucrose g 28.8 | Sucrose g 28.8 |
| Salt Mix g 10.8 | Salt Mix g 10.8 |
| Torula Yeast g 14.4 | Torula Yeast g 14.4 |
| **Cellulose g 0** | **Cellulose g 21.6** |
| Cholesterol g 3.15 | Cholesterol g 3.15 |
| Sorbic Acid g 1.8 | Sorbic Acid g 1.8 |
| Ascorbic Acid g 3.6 | Ascorbic Acid g 3.6 |
| VanderZandt Vit Mix g 12.6 | VanderZandt Vit Mix g 12.6 |
| Water (cool) ml 450 | Water (cool) ml 450 |
| Water (boiling) ml 450 | Water (boiling) ml 450 |
| Linseed oil ml 6 | Linseed oil ml 6 |

**Important note:**

The casein used is Teklad Vitamin-Free. In this diet, casein is the predominant source of protein. If another type of casein is used, the amount of vitamins may also vary.

**2. Is hindwing melanisation continuous or discrete?**

To evaluate the extent to which human-eye categorization of moth melanin morphs (high melanin and low melanin) captures variation in the degree of melanisation, we compared human-eye categories to the results from a cluster analysis of the proportion of the wing that was melanised. We used k-means clustering with $k = 2$ to sort melanisation measurements into two clusters. The resulting clusters were labelled high and low melanin clusters based on their mean melanisation scores (e.g., the cluster with the higher mean melanisation score was labelled the high melanin cluster). We then calculated the proportion of moths that were "correctly" classified – those for which the human-eye and cluster analysis scores were the same – as: [(human-eye high melanin moths clustered as high melanin) + (human-eye low melanin moths clustered as low melanin)] / (total number of moths).

A low correspondence between human-eye categorizations and cluster analysis results does not necessarily indicate that moth melanisation is best treated as a continuous variable; it is still possible that the proportion of the wing that is melanised is well captured in a small number of categories different from the high and low categories used previously (e.g., high, medium, and low; or high and low but with a different threshold). To test this possibility, we determined the optimal number of clusters in the proportion of the wing that is melanised using the silhouette method (Rousseeuw, 1987). This approach calculates a silhouette score for each tested k value, with higher silhouette scores indicating better support for that k value. The k value with the highest silhouette score best captures the structuring of the data (the clusters are maximally distinguishable from one another), and is therefore considered the optimal number of clusters. We calculated silhouette scores for k = 2 through k = 10 (i.e., sorting proportion melanised into two to ten clusters).

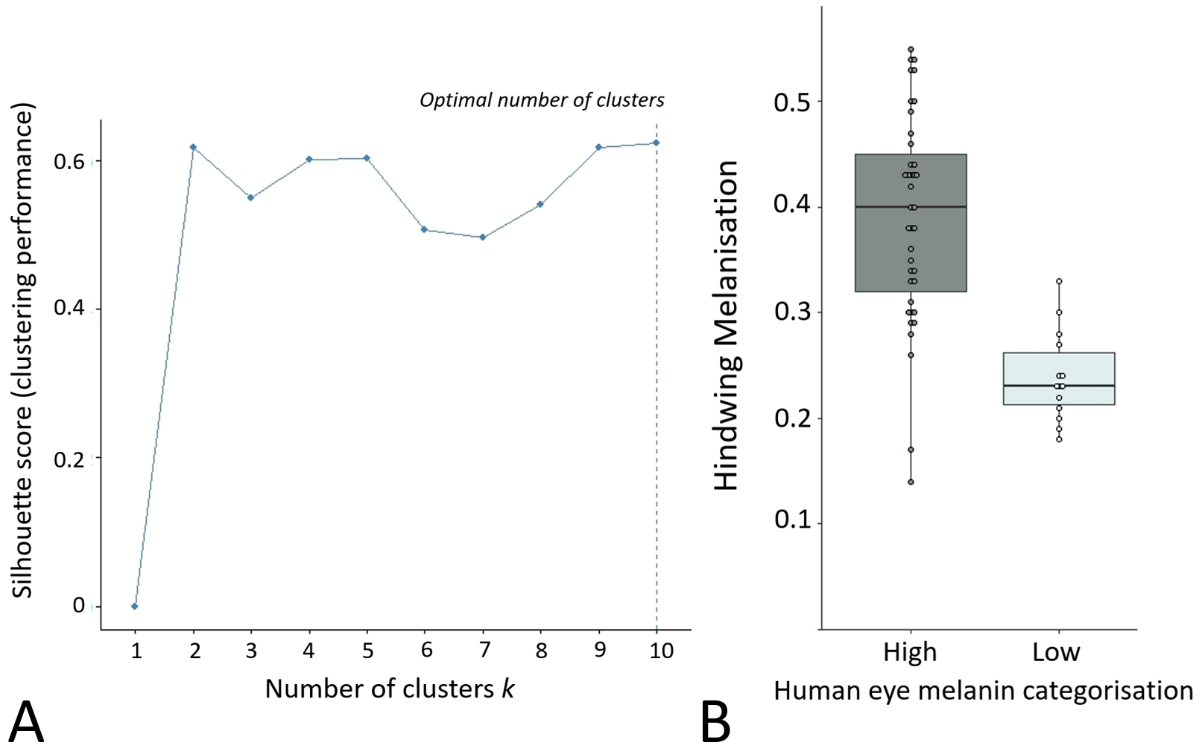


**Figure 1 A)** The k value with the highest silhouette score best captures the structuring of the data (optimal number of clusters). While an unimodal plot would have indicated a clearly optimal k value, this plot suggests more continuous data in the hindwing melanin amount. **1 B)** Objective hindwing melanisation measurements (the proportion of the wing that is melanised) for high- and low-melanin morphs as scored by the human eye. Note the overlap between high and low melanisation human-generated scores: subjective scoring may not be sensitive enough to detect patterns of variation.

When the proportion of the wing that was melanised was clustered into two categories, 70% of the resulting moth classifications corresponded with categorizations based on the human eye. To identify the optimal number of melanin clusters k we used the silhouette score, which represented the pairwise dissimilarity between and within cluster distances (clustering performance). The proportion of the wing that was melanised was best represented by 10 clusters, although other k values had very similar degrees of support (Fig. 1A); this high ‘optimal’ k value and the similar degrees of support for multiple different k values indicates that wing melanisation is best treated as a continuous variable. Therefore, in the analyses we considered the melanin amount in the wings as continuous.

### 3. Effect of diet manipulation and hindwing melanin on the predator response to male's defensive fluid

#### 3.1 Latency to approach

**Table S1 A)** Latency to approach the bait (oat) with control (water) was tested using cox mixed-effects model using package coxme, fit by maximum likelihood. The latency to approach was set as response variable; the different types of diet (low protein, high protein) were set as predictor variables. The diet and trial were set as fixed effects, while bird ID was set as a random effect. **S1 B)** Latency to approach the bait (oat) without control was tested using cox mixed-effects model using package coxme, fit by maximum likelihood. The latency to approach was set as response variable; the different types of diet were set as predictor variables. The interaction between diet (low protein, high protein), the amount of melanin and trial were set as fixed effects, while bird ID was set as a random effect.

| S1A) With control | coeff | exp(coef) | se(coef) | z | p |
|---|---|---|---|---|---|
| **High prot. treatment** | -0.49 | 0.61 | 0.58 | -0.85 | 0.39 |
| **Low prot. treatment** | -0.22 | 0.80 | 0.57 | -0.39 | 0.70 |
| **trial 3** | 0.29 | 1.34 | 0.25 | 1.14 | 0.25 |

| S1B) Without control | coeff | exp(coef) | se(coef) | z | p |
|---|---|---|---|---|---|
| **Low prot. treatment** | 0.34 | 1.41 | 1.25 | 0.27 | 0.78 |
| **Melanin hindwing** | -3.52 | 0.03 | 2.41 | -1.46 | 0.14 |
| **trial 3** | 0.17 | 1.19 | 0.22 | 0.78 | 0.44 |
| **Low protein diet:Hindwing melanin** | 1.02 | 2.78 | 3.16 | 0.32 | 0.75 |

**Spearman's rank correlation** (Correlation tested between the hindwing melanin and the predators' response (latency to approach) to the chemical defence) $r_s = 0.22$; $p = 0.265$

### 3.2 Latency to eat after approaching

**Table S2 A)** Latency to eat after approaching the bait (oat) with control (water) was tested using cox mixed-effects model using package coxme, fit by maximum likelihood.The latency to eat after approaching was set as response variable; the different types of diet (low protein, high protein) were set as predictor variables. The diet and trial were set as fixed effects, while bird ID was set as a random effect. **S2 B)** Latency to eat after approaching the bait (oat) without control was tested using cox mixed-effects model using package coxme, fit by maximum likelihood. The latency to eat after approaching was set as response variable; the different types of diet were set as predictor variables. The interaction between diet (low protein, high protein), the amount of melanin and trial were set as fixed effects, while bird ID was set as a random effect.

| S2 A) With control | **coeff** | **exp(coef)** | **se(coef)** | **z** | **p** |
|---|---|---|---|---|---|
| **High prot. treatment** | 0.27 | 1.32 | 0.34 | 0.80 | 0.42 |
| **Low prot. treatment** | -0.15 | 0.86 | 0.34 | -0.44 | 0.66 |
| **trial 3** | -0.12 | 0.89 | 0.24 | -0.48 | 0.63 |

| S2 B) Without control | **coeff** | **exp(coef)** | **se(coef)** | **z** | **p** |
|---|---|---|---|---|---|
| **Low prot. treatment** | -0.06 | 0.94 | 0.25 | -0.25 | 0.80 |
| **Melanin hindwing** | -0.67 | 0.51 | 0.96 | -0.70 | 0.48 |
| **trial 3** | 0.08 | 1.08 | 0.22 | 0.37 | 0.71 |

**Spearman's rank correlation**(Correlation tested between the hindwing melanin and the predators' response (latency to eat after approaching) to the chemical defence) $r_s$ = 0.039; *p = 0.71*

### 3.3 Latency to eat

**Table S3** Latency to eat the bait (oat) with control (water) was tested using cox mixed-effects model using package coxme, fit by maximum likelihood. The latency to eat was set as response variable; the different types of diet (low protein, high protein) were set as predictor variables. The diet and trial were set as fixed effects, while bird ID was set as a random effect.

| S3 With control | **coeff** | **exp(coef)** | **se(coef)** | **z** | **p** |
|---|---|---|---|---|---|
| **High prot. treatment** | 0.27 | 0.76 | 0.53 | -0.52 | 0.60 |
| **Low prot. treatment** | -0.42 | 0.65 | 0.52 | -0.81 | 0.42 |
| **trial 3** | -0.02 | 0.98 | 0.26 | -0.07 | 0.94 |

### 3.4 Beak wiping

**Table S4 A)** Beak wiping reaction to the bait (oat) with control (water) was tested using a generalised linear mixed-effects models (GLMM) with a log link and Poisson distribution, fit by maximum likelihood (Laplace approximation) to test for differences in beak wiping events per minute. The diet (low protein, high protein) and trial were set as fixed effects, while the beak wiping frequency was set as response variable and bird ID as a random effect. **S4 B)** Beak wiping reaction to the bait (oat) without control was tested using a generalised linear mixed-effects models (GLMM) with a log link and Poisson distribution, fit by maximum likelihood (Laplace approximation) to test for differences in beak wiping events per minute. The interaction between diet (low protein, high protein), the amount of melanin, and trial were set as fixed effects, while the beak wiping frequency was set as response variable and bird ID as a random effect

| S4 A) With control | **Estimate** | **Std. Error** | **z value** | **Pr(>\|z\|)** |
|---|---|---|---|---|

| | | | | |
|---|---|---|---|---|
| **Intercept** | -4.41 | 0.70 | -6.34 | 2.28e-10 *** |
| **High prot. treatment** | 0.09 | 0.75 | 0.12 | 0.90 |
| **Low prot. treatment** | 0.03 | 0.76 | 0.04 | 0.97 |
| **trial 3** | 0.06 | 0.25 | 0.22 | 0.82 |

| S4 B) Without control | **Estimate** | **Std. Error** | **z value** | **Pr(>\|z\|)** |
|---|---|---|---|---|
| **Intercept** | -3.53 | 0.87 | -4.03 | 5.49e-05 *** |
| **Treatment** | -0.72 | 1.19 | -0.60 | 0.55 |
| **Hindwing melanin** | -3.40 | 2.27 | -1.49 | 0.13 |
| **trial 3** | 0.008 | 0.20 | 0.04 | 0.97 |
| **Low protein diet:Hindwing melanin** | 3.13 | 2.95 | 1.06 | 0.29 |

**Spearman's rank correlation** (Correlation tested between the hindwing melanin and the predators' response (beak wiping) to the chemical defence) $r_s$ = -0.025; $p = 0.8$
